# ADAM17 is the main sheddase for the generation of human triggering receptor expressed in myeloid cells (hTREM2) ectodomain and cleaves TREM2 after histidine 157

**DOI:** 10.1101/133751

**Authors:** Dominik Feuerbach, Patrick Schindler, Carmen Barske, Stefanie Joller, Edwige Beng-Louka, Katie A Worringer, Sravya Kommineni, Ajamete Kaykas, Daniel J Ho, Chaoyang Ye, Karl Welzenbach, Gaelle Elain, Laurent Klein, Irena Brzak, Anis K Mir, Christopher J Farady, Reiner Aichholz, Simone Popp, Nathalie George, Christine L Hsieh, Mary C Nakamura, Ulf Neumann

## Abstract

Triggering receptor expressed in myeloid cells (TREM2) is a member of the immunoglobulin superfamily and is expressed in macrophages, dendritic cells, microglia, and osteoclasts. TREM2 plays a role in phagocytosis, regulates release of cytokine, contributes to microglia maintenance, and its ectodomain is shed from the cell surface. Using both pharmacological and genetic approaches we report here that the main protease contributing to the release of TREM2 ectodomain is ADAM17, (a disintegrin and metalloproteinase domain containing protein, also called TACE, TNFα converting enzyme) while ADAM10 plays a minor role. Using mutational analysis, we demonstrate that the main cleavage site of the sheddases is located within the stalk region of TREM2 proximal to the plasma membrane. Complementary biochemical experiments reveal that cleavage occurs between histidine 157 and serine 158. Shedding is not altered for the R47H-mutated TREM2 protein that confers an increased risk for the development of Alzheimers disease. O-glycosylation is detected within the stalk region, but distant to the cleavage site. These findings reveal a link between shedding of TREM2 and its regulation during inflammatory conditions or chronic neurodegenerative disease like AD in which activity or expression of sheddases might be altered.

## Introduction

Triggering receptor expressed in myeloid cells (TREM2) is a type I transmembrane glycoprotein and a member of the immunoglobulin (Ig) receptor superfamily (1). TREM2 expression has been shown in macrophages, dendritic cells, microglia and osteoclasts (2-4), and expression seems to be temporally and spatially regulated. In macrophages expression is upregulated during the course of inflammation, e.g. expression peaks 2-3 days after thioglycolate challenge in a murine model of peritonitis (5). TREM2 is also enriched at those microglia cell surface regions which contact Aβ plaques or neuronal debris (6). Some of the ligands that are sensed by TREM2 in this environment have recently been identified, for example phospholipids and myelin lipids (7) as well as ApoE (8,9). Other ligands could be Aβ and plaque associated neuronal debris since TREM2 contributes to the uptake of Aβ into microglia (10).

This is well in line with earlier genome wide association studies showing that a TREM2 SNP, rs75932628-T encoding the R47H variant, conferred a significantly increased risk of late onset Alzheimer`s disease (LOAD) with odds ratios of 5.05 (11) and 2.92 (12). These odds ratios are comparable with that of the well-established AD risk gene APOE4 (13). It is of note that R47H mutated TREM2 displays almost the same cell surface expression as WT TREM2 (14) but is functionally impaired: the R47H mutation in TREM2 reduces binding of lipid ligands (15) and ApoE (8,9). The mutation also reduces phagocytic capacity (14) and impedes recycling of TREM2 via Vps35 within the retromer complex (16). Some human R47H carriers without AD have been characterized, and these individuals lose brain volume faster than non-carriers (17), have a poorer cognitive function than age matched controls, (12) and show upregulation of pro-inflammatory cytokines (e.g. RANTES, INFγ) and downregulation of protective markers (e.g. IL-4, ApoA1; (18)).

The shed ectodomain of TREM2 (sTREM2) in human CSF has been assessed as a potential AD biomarker and has been shown to be increased during ageing in general (19). Detailed analysis during the course of AD revealed that sTREM2 increases early in AD before clinical symptoms appear, peaks in MCI-AD, and stays elevated but at lower levels compared to the MCI-AD stage in AD dementia (19).

sTREM2 comprises the IGSF domain and part of the stalk region, but the exact size is a matter of debate. The entire TREM2 protein consists of a leading signal peptide (amino acids 1–13), a single V-type IgSF extracellular region, (95 AA / 14-113), a stalk region (amino acids 114–174 / 64 AA), a positively charged transmembrane domain (amino acids 175–197 / 24 AA), and a cytosolic tail (amino acids 198–230 / 32 AA) (1). In spite of the long cytosolic tail, there is no signaling motif. Instead, TREM2 forms a heterodimer with DAP12 and the lysine within the transmembrane region of DAP12 interacts with aspartic acid in the transmembrane part of TREM2 (1). Shedding of TREM2 is thought to involve as first step ADAM10 (a disintegrin and metalloproteinase domain containing protein) and/or ADAM17 (also termed TACE, TNFα converting enzyme) (14). This proteolytic cleavage liberates sTREM2 from the plasma membrane. Next, γ-secretase cuts the membrane associated C-terminal fragment (CTF) enabling further degradation of the peptide (20). Inhibition of γ-secretase results in accumulation of CTF and trapping of DAP12 within this complex. This leads to reduced TREM2 signaling as availability of DAP12 becomes limiting.

In the current study we have investigated the differential contribution of ADAM10 and ADAM17 to shedding of TREM2 ectodomain using a pharmacological and a genetic approach. Next we identified the cleavage site of ADAM10/ADAM17 within the stalk region of TREM2 using a mutational and complementary biochemical approaches.

## Results

### ADAM17 inhibitors stabilize TREM2 at the cell surface

To determine the contribution of ADAM10 or ADAM17 to shedding of TREM2 ectodomain we first set out to find selective inhibitors and identified two compounds (DPC333 (21) and GI254023 (22)) that were characterized for inhibitory selectivity towards ADAM10 and ADAM17 (see supplementary figure 1). While DPC333 is a more potent inhibitor on ADAM17 (IC_50_ < 0.6 nM) than on ADAM10 (IC_50_= 5.3 nM), GI254023 displays selectivity for ADAM10 (IC_50_ 1.5 nM) over ADAM17 (IC_50_= 196 nM). Using live cell imaging, cell surface expression of hTREM2 was assessed in CHO-hDAP12-hTREM2 cells after overnight treatment of the cells with the two ADAM inhibitors under conditional shedding conditions (figure 1A) or after treatment of the cells with PMA (figure 1B). The ADAM17 selective inhibitor DPC333 dose-dependently increases TREM2 cell surface levels under both conditions. A limited effect on TREM2 cell surface levels is also observed at higher concentrations for GI254023, but only under steady state conditions. This effect might be attributed to true ADAM10 inhibition or it might be caused by unspecific inhibition of ADAM17 by GI254023 when used at high concentrations. In PMA-treated cells there is a complete lack of effect of GI254023 on TREM2 cell surface expression (figure 1B). To get closer to a physiological cellular system, a similar experiment was conducted in human M2A macrophages differentiated from CD14^+^ human monocytes (figure 1 C and D). These results replicate very well the initial findings in CHO-hDAP12-hTREM2 cells; the ADAM17 inhibitor DPC333 increases TREM2 cell surface expression dose-dependently under both conditions (figure 1C and 1D) and the selective ADAM10 inhibitor displays a small effect on steady-state shedding (figure 1C). In summary, these experiments indicate that in human macrophages, ADAM17 plays a critical role for TREM2 shedding but a marginal contribution of ADAM10 under steady state condition cannot be excluded.

**FIGURE 1.**
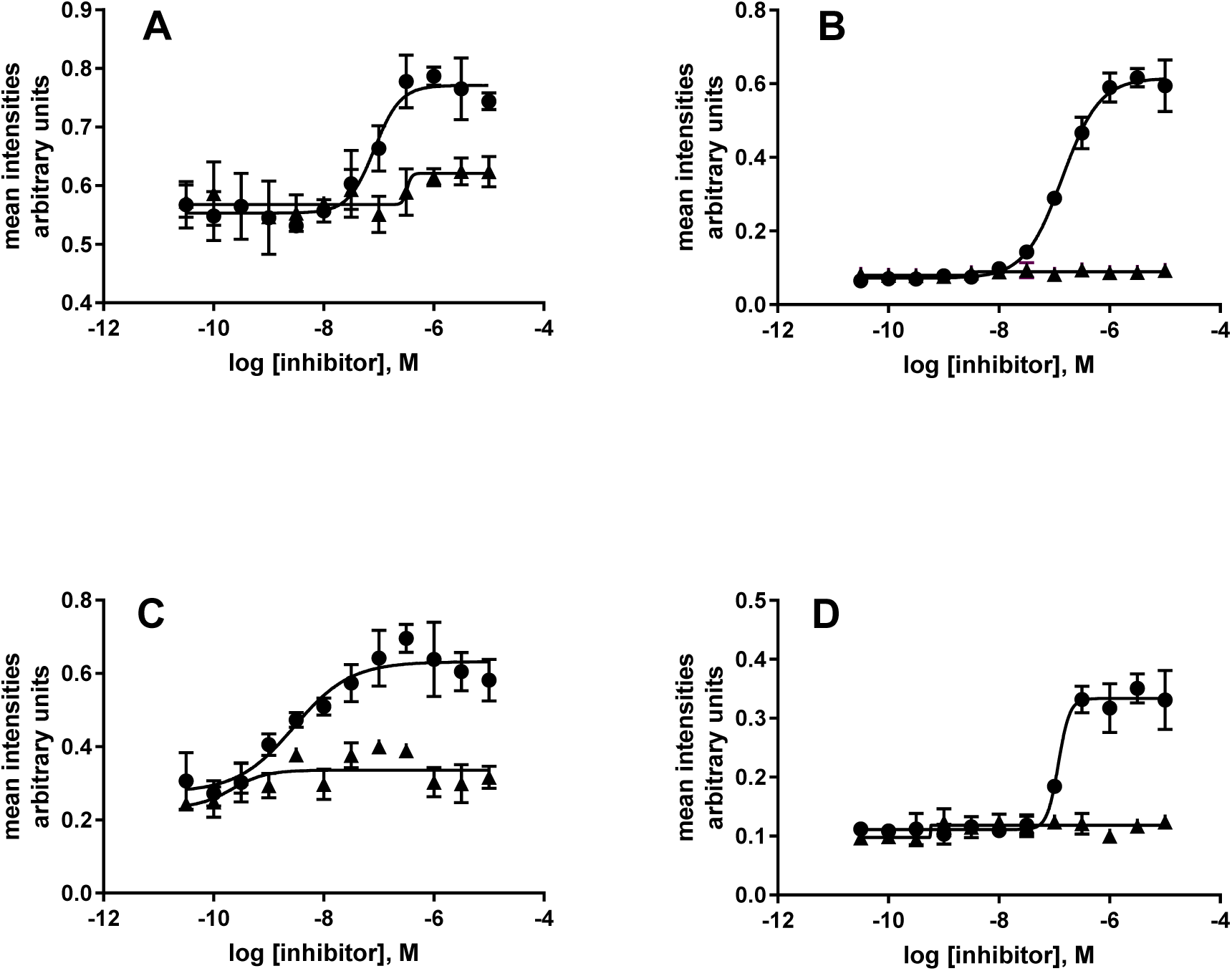
ADAM17 is the pivotal sheddase for the cleavage of TREM2 ectodomain in CHO-hDAP12-hTREM2 cells and human M2A macrophages. Cell imaging of TREM2 cell surface expression in CHO-hDAP12-hTREM2 (A and B) or human M2A macrophages (C and D) after treatment with ADAM inhibitors DPC333 (black circle) or GI254023 (upward triangle). Additional PMA treatment was applied in panel B and D. TREM2 cell surface staining is plotted as mean intensities corrected for nuclear staining as mean ± S.E. (n=3). A representative experiment from two experiments is shown.

### ADAM17 ablation in THP1 cells reduces constitutive shedding

To confirm these conclusions, a genetic approach was used to further investigate the contribution of ADAM10/17 to TREM2 shedding. Human monocytic THP1 cells were chosen as model system which endogenously expresses TREM2. Employing the CRISPR/CAS9 technology, clones were generated that lack expression of ADAM10 (AD10 H4) or ADAM17 (AD17 G12) as well as a control cell line (Ctrl gRNA). Absence of gene products was verified by FACS analysis or Western blot (see supplementary figure 2 and 3). Cell surface expressed TREM2 and sTREM2 were assessed in the three cell lines in the same experiment using three different conditions: no treatment reflecting constitutive shedding, PMA treatment that maximally activates shedding, and finally PMA and DPC333 treatment. Loss of ADAM17 increases TREM2 cell surface expression and strongly reduces soluble TREM2 under conditional shedding conditions, whereas lack of ADAM10 has no significant effects (see black bars in figure 2A and B), thus indicating that ADAM17 is the main sheddase contributing to constitutive shedding. Maximally activating sheddases with PMA leads to a strong reduction of cell surface TREM2 both in the control cell line and in the ADAM10 deficient clone. This is reflected in a strong increase in sTREM2. In the ADAM17 deficient cell line, PMA treatment also leads to a reduction of cell surface TREM2, but to a smaller extent than in the control CRISPR clone. Also, the increase in sTREM2 is smaller compared to PMA treated control CRISPR clone and ADAM10 deficient clone. However, the cleavage of TREM2 in the AD17 G12 clone in the presence of PMA must be caused by a sheddase other than ADAM17. Accordingly, in the ADAM10 H4 clone the increase in PMA induced shedding might be caused by additional activation of ADAM17. Co-treatment with PMA and DPC333 restored TREM2 cell surface levels in Ctrl gRNA and AD10 H4 cells but had smaller effects in the AD17 G12 clone. In the AD17 G12 clone cell surface TREM2 levels do not reach the extent that is seen under constitutive shedding conditions. sTREM2 however is strongly reduced in the AD17 G12 clone under these conditions compared to PMA treatment only. In summary, in THP1 cells ADAM17 seems to be the main sheddase responsible for constitutive shedding. After PMA treatment additional shedding mechanisms come into play, one of which might involve ADAM10.

**FIGURE 2.**
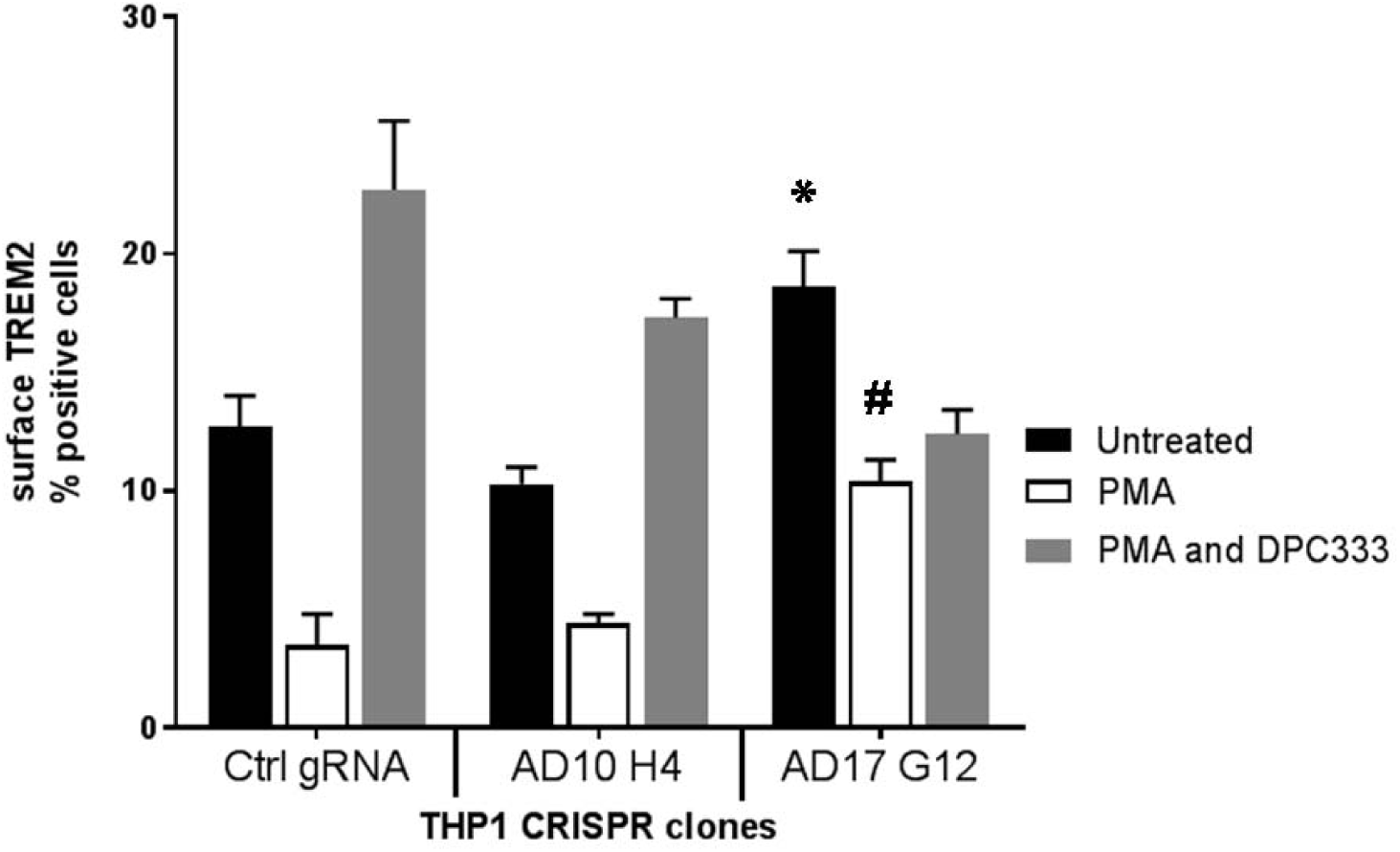
ADAM17 but not ADAM10 TREM2 CRISPR knockout THP1 cells show increased TREM2 expression and reduction of sTREM2. A, FACS analysis of TREM2 cell surface expression in THP1 CRISPR cell clones. Data are plotted as mean ± S.E. (n=2). * p < 0.01, statistical difference to untreated group of Ctrl gRNA clone. # p < 0.01, statistical difference to PMA treated group of Ctrl gRNA clone.

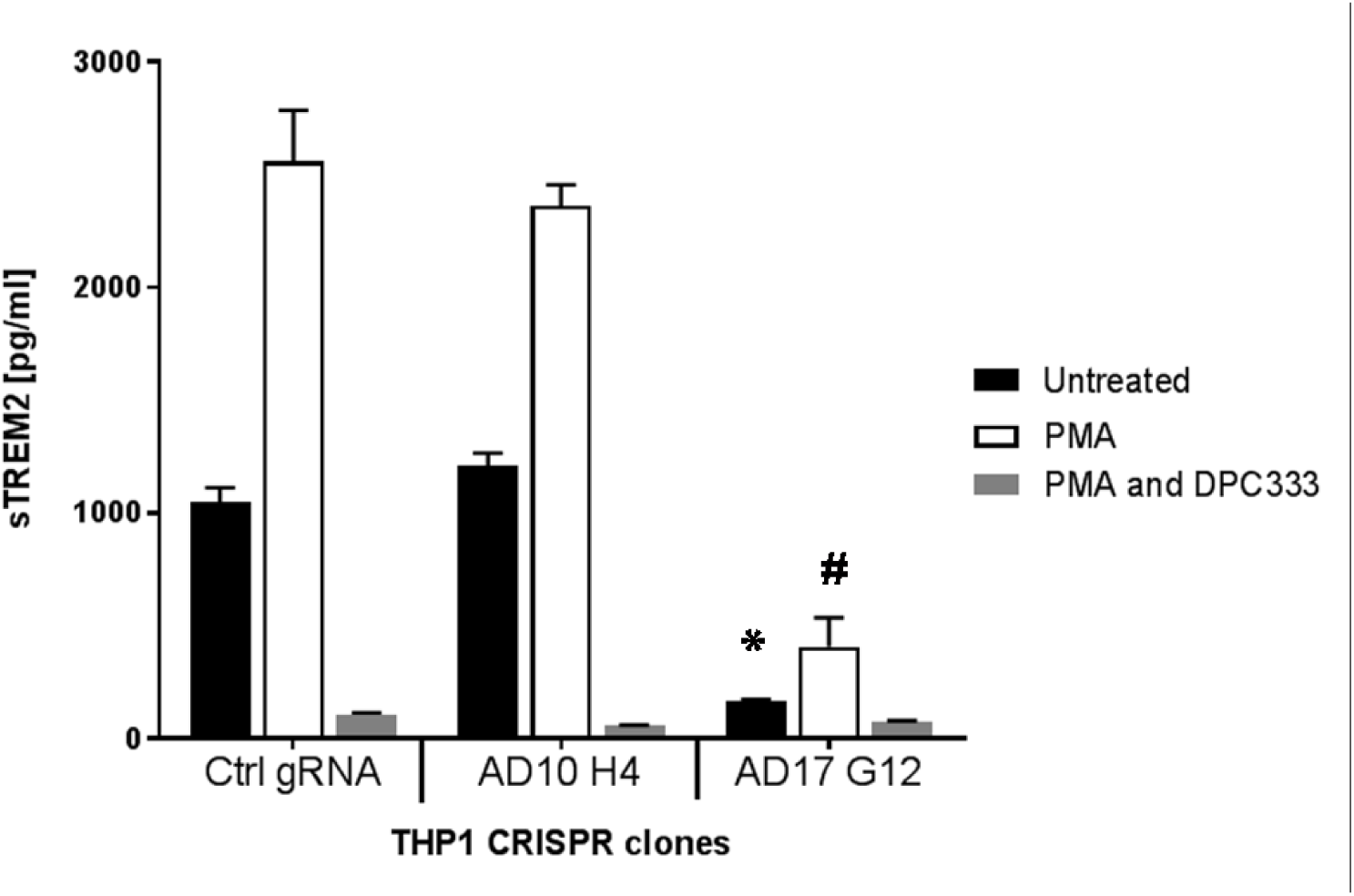
B. sTREM2 analysis of supernatants from THP1 C CRISPR cell clones. * p < 0.01, statistical difference to untreated group of Ctrl gRNA clone. # p < 0.01, statistical difference to PMA treated group of Ctrl gRNA clone. CtrlgRNA, control gRNA transfected clone; AD10 H4, ADAM10 CRISPR knockout clone; AD17 G12 ADAM17 CRISPR knockout clone. Data are plotted as mean ± S.E. (n=2).

### 2 AA stretches close to the TM are important for shedding

In the next experiments we used site directed mutagenesis to identify areas within TREM2 stalk region that harbor the cleavage site or are important for binding of the sheddases. WT TREM2 or the respective mutants were transfected together with hDAP12 into HEK-FT cells. 48 h later cells were treated for 30 min with PMA to activate ADAMs at cell surface (23). TREM2 cell surface expression was assessed by FACS, and results are presented as expression ratio of untreated over PMA treatment (figure 3A). The evaluation of each construct in the presence and absence of PMA treatment overcomes the possibility that some constructs might display different binding properties for the antiserum and allows direct comparison of changes in TREM2 cell surface expression after activation of sheddases. A ratio of 1 indicates complete inhibition of shedding. An overview of different mutations that have been generated is shown in figure 3B. Deletion of the first 16 AA proximal to the TM maximally reduces TREM2 shedding (TRUNCIII-159-174). This suggests that this region might entail the cleavage and/or the binding site for ADAMs. Next we generated 4 shorter deletion mutants encompassing this area, each 6 AA long (TRUNC1, T2del3-8, T2del6-11 and T2del11-16). While there was no or very little effect on shedding for mutants TRUNC1, T2del3-8 and T2del6-11, the mutant T2del11-16 showed reduced PMA induced shedding. We then designed three AA-replacements to overcome the issue that deletion mutants might shift the cleavage site closer to the transmembrane region causing reduction of cleavage due to steric hindrance. We replaced AA 156-164 and 169-172 with larger hydrophobic residues that should render the stalk region resistant to protease cleavage (24). While T2-YGGWGGWGP showed a trend for reduction of TREM2 shedding, replacement of 4 membrane-proximal AA was more efficacious (T2-WFRW) and the combination of both (T2-double) had a similar effect as the deletion mutant TRUNCIII-159-174. In summary the mutagenesis approach revealed that two areas are important for PMA induced shedding of TREM2, a membrane proximal at AA 169-172 and a membrane distal in the region AA 156-164.

**FIGURE 3.**
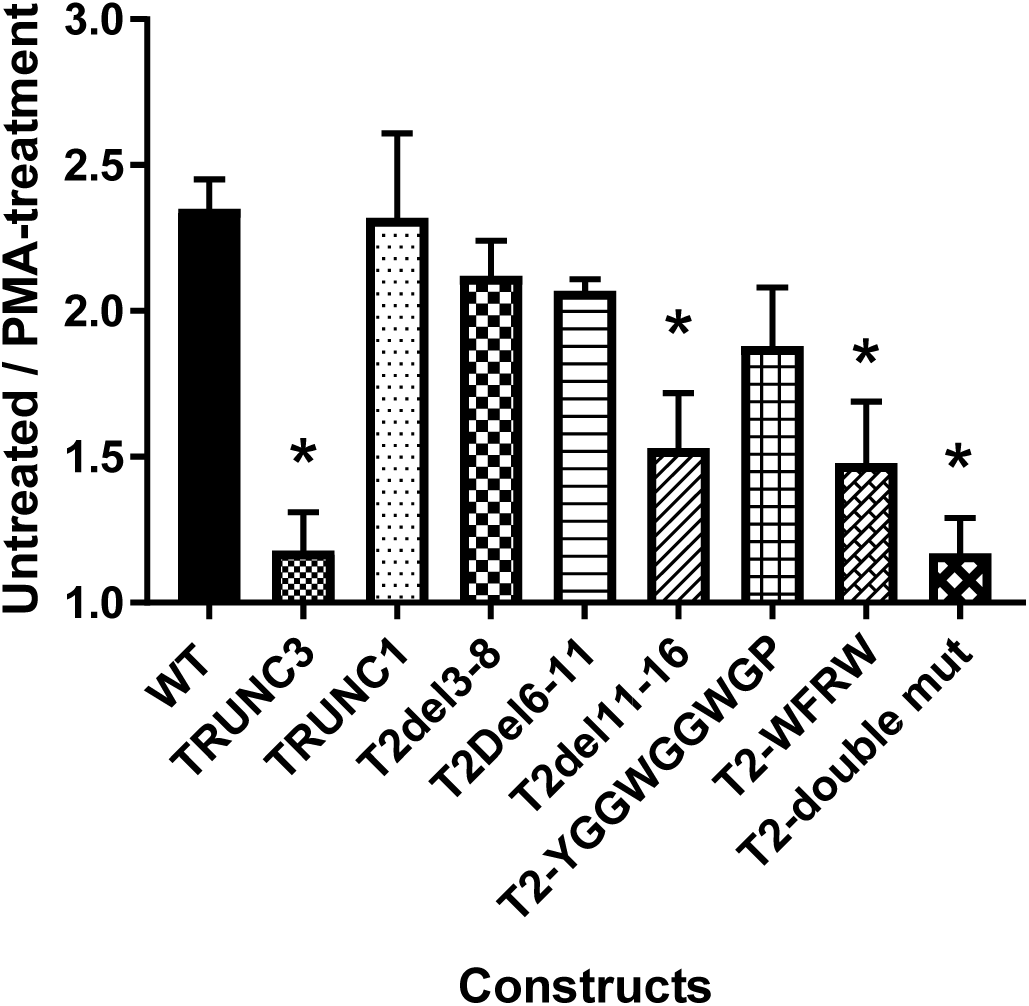
Two AA stretches in the membrane proximal part of the TREM2 stalk region are important for shedding. A, FACS analysis of TREM2 WT and mutants transiently expressed in HEK293-FT cells. Cells were treated 48 h after transfection with 50 ng/ml PMA or 0.05% DMSO. Cells were detached, labelled with AF1828 antiserum and analyzed by flow cytometry. Data are plotted as the ratio of untreated over PMA treatment for each mutant as mean ± S.E. (n=3). Statistical differences to WT were calculated by Anova with Dunnett’s multiple comparison test. *P < 0.01.

**Figure 3B.**
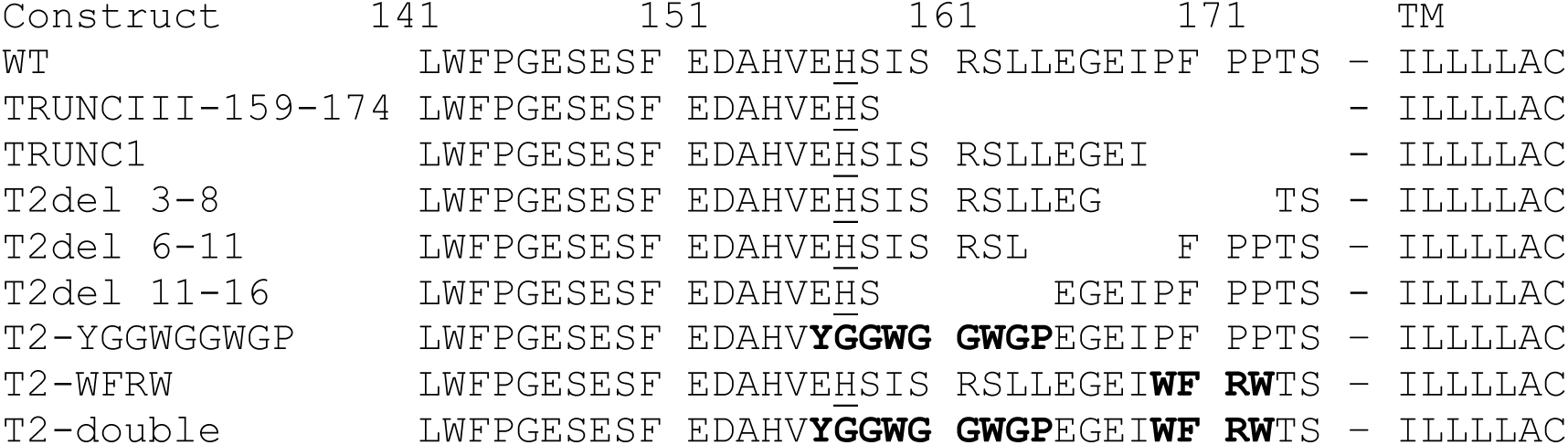
B, Amino acid representation of hTREM2 mutants. AA numbering according to iProt Q9NZC2. TM: transmembrane region. Gaps indicate deleted AA within the respective mutant. Exchanged AA in bold. Underlined AA indicates main sheddase cleavage site for generation of the C-terminus of TREM2 ectodomain as identified in figures 4-6.

Next we assessed whether the T2-double-TREM2 construct retained functionality. To this end this mutant was stably transfected into BWZ cells that already expressed mouse DAP12 and a beta-Gal reporter driven by NFAT (25). mDAP12 expressed without TREM2 in this cell lines represents background reporter gene activity (RGA) in this system. Comparing RGA after TREM2 activation in BWZ-T2-double and BWZ-TREM2-WT cells revealed comparable activity, indicating that mutated T2-double is still capable of signaling via DAP12 (figure 3C).

**Figure 3C.**
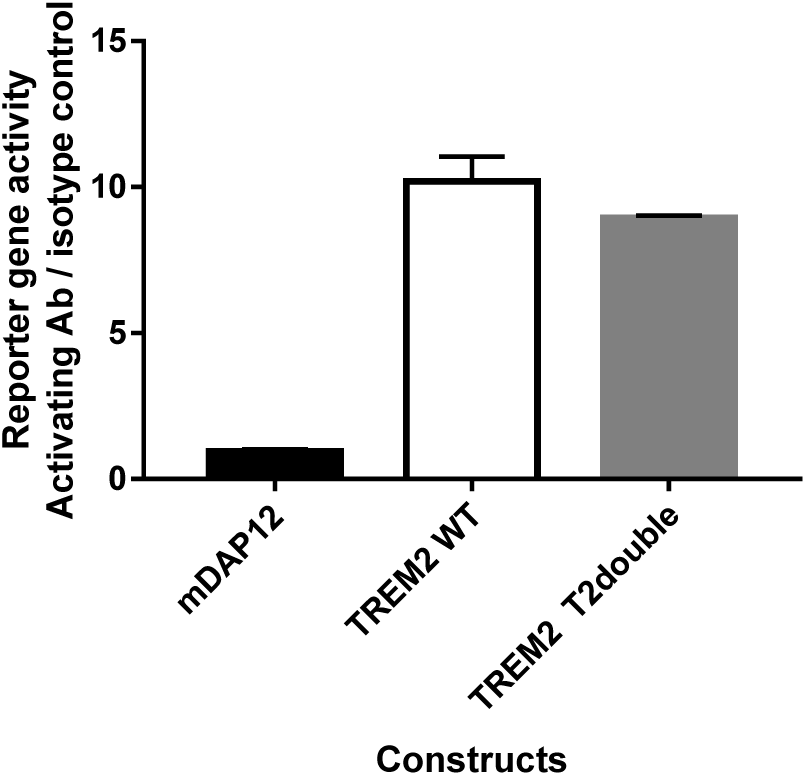
C, TREM2 double construct was stably expressed in BWZ-lacZ-mDAP12 cells. Cells were seeded on activating Ab MAB17291 or isotype control and reporter gene activity assessed after 16 h. Data are plotted as the ratio of RGA for activating/control Ab as mean ± S.E. (n=3).

### ADAM17 cleaves stalk region peptides at position H157-S158

In order to unambiguously identify the cleavage site of ADAM10/17 within the stalk region of TREM2, we investigated *in vitro* cleavage patterns of stalk region-derived peptides and confirmed the cleavage site by determination of the C-terminus of shed soluble TREM2 from cell culture supernatants. We first designed a series of peptides covering the stalk region for *in vitro* investigation of ADAM10/17 cleavage (Fig 4A). All peptides were obtained with an N-terminal 7-methoxycoumarin (Mca) fluorescent tag. Peptides 1-3, 1a and 2a were incubated with ADAM17 in neutral buffer for up to 48 hours, and reaction was followed by HPLC analysis of aliquots withdrawn at different time points. Significant cleavage was observed only for peptide 3, while peptides 1, 2, 1a, and 2a showed only very little or no reduction of the parent peptide (Fig 4B). HPLC-MS analysis of the reaction mixture of peptide3/ADAM17 identified one major product, SISRSLLEGEIPFP-NH_2_, together with 2 minor cleavage products (Fig 4B). This suggests that the H157-S158 bond in TREM2 is the main cleavage site for ADAM17. HPLC analysis of the incubation mixture at times > 24 hours showed appearance of more than one product peak, together with a reduced main product peak. At these time points there was essentially no substrate left. Therefore we conclude that the minor cleavage products originate from secondary ADAM17 cleavage of the main product. The same analysis was carried out with ADAM10, and a very similar cleavage pattern was obtained (data not shown).

**FIGURE 4A.**
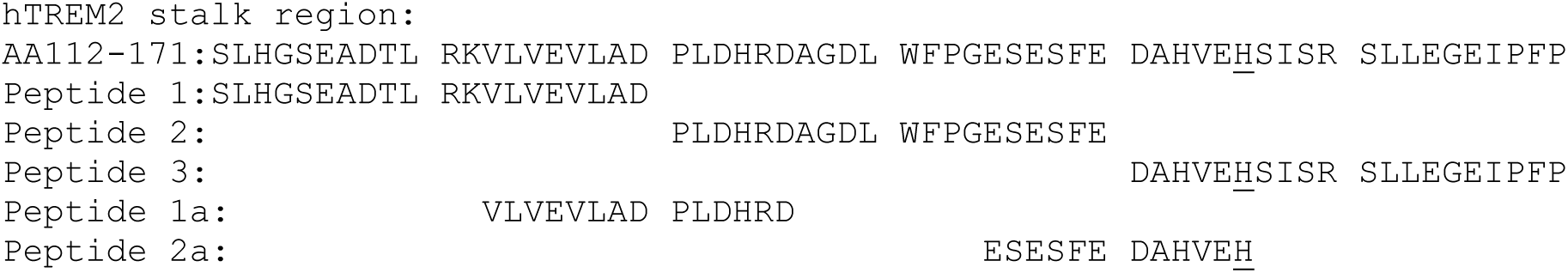
ADAM17 cleaves TREM2 stalk region peptides *in vitro* at H157-S158. A, Stalk region derived synthetic peptides for *in vitro* cleavage assays. All peptides were obtained with an N-terminal 7-methoxycoumarin (Mca) fluorescent tag at the C-terminus. Underlined AA indicates main sheddase cleavage site.

**FIGURE 4B.**
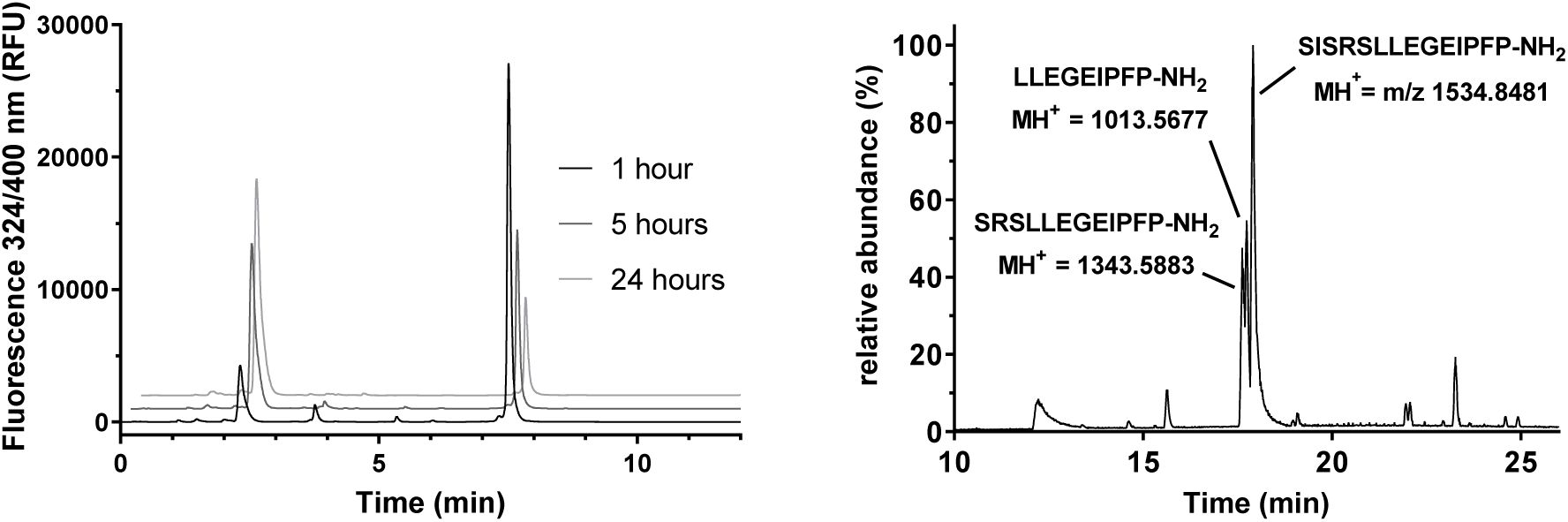
B, HPLC analysis of cleavage of peptide 3 (10 μM) by ADAM17 (31 nM). B HPLC-MS analysis of incubation mixture of peptide 3 with ADAM17 for 48 hours, with identification of the major product and 2 minor products. C Time course of ADAM17 cleavage of peptides 1, 2, and 3 (mean of 2 experiments).

### TREM2 ectodomain shed from cells is cleaved at H157

To substantiate the *in vitro* findings from the peptide analysis on sheddase cleavage site of TREM2 in a cellular system, HEK-FT cells were transiently transfected with hTREM2 or hTREM2-R47H in combination with hDAP12. Transfected cells from both conditions were treated with PMA or solvent. sTREM2 was immunopurified from cellular supernatant and subjected to trypsin or Asp-/Glu-C enzyme digestion followed by analysis of the peptides by LC-MS. In all four conditions the same N-terminal peptide was identified (D_137_-H_157_) indicating as main cleavage site H157 (figure 5 and figure S4). These experiments show that neither PMA treatment nor the R47H mutation induce a shift of the main cleavage site.

**FIGURE 5.**
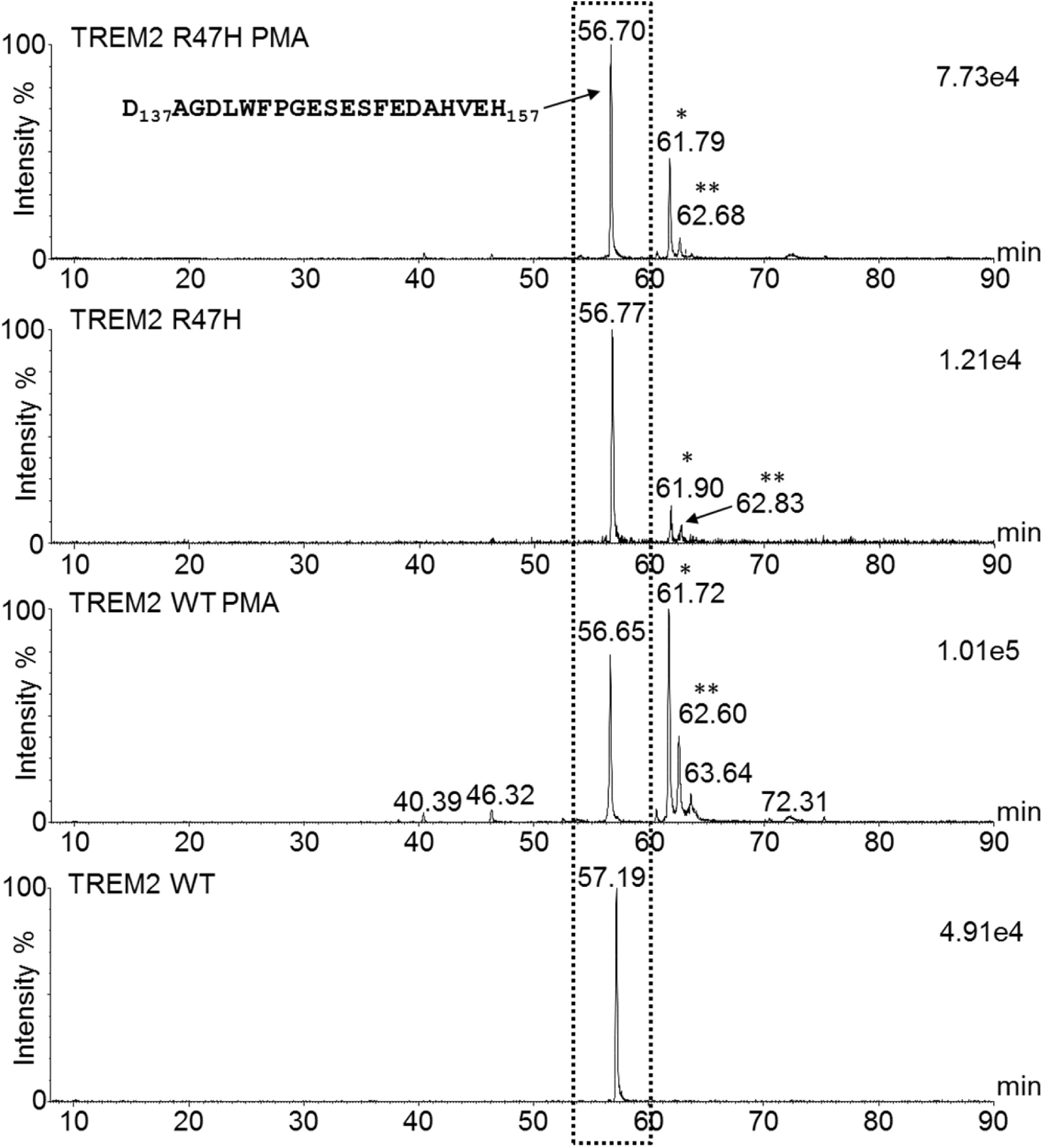
Identification of C-terminus of TREM2 ectodomain shed from HEK-FT cells transiently transfected with WT or R47H human TREM2 and DAP12. Ion extracts of the 3-time charged [D137-H157] peptide ion, m/z 791.94-792.06. The peptide ion of interest is clearly present in all four shed TREM2 tryptic digests (boxed ion extract). * represents an unknown peptide present as a 5-time charged ion and ** is the deaminated form of the same peptide.

In the next experiments we set out to identify the entire shed TREM2 ectodomain from the immunopurified cellular supernatant treated with PNGase-F and sialidase-A followed by LC-MS analysis. We were able to detect a 15,619 Dalton peptide species in the supernatant from cells transfected with WT TREM2 which corresponds to de-glycosylated TREM2_19-157_ (figure 6). A corresponding peptide of 15,600 Daltons was detected in the immunopurified cellular supernatant of R47H-TREM2 transfected cells. Due to the AA exchange this peptide is 19 Daltons lighter than the WT peptide and confirms the same cleavage site for mutated R47H-TREM2.

**FIGURE 6.**
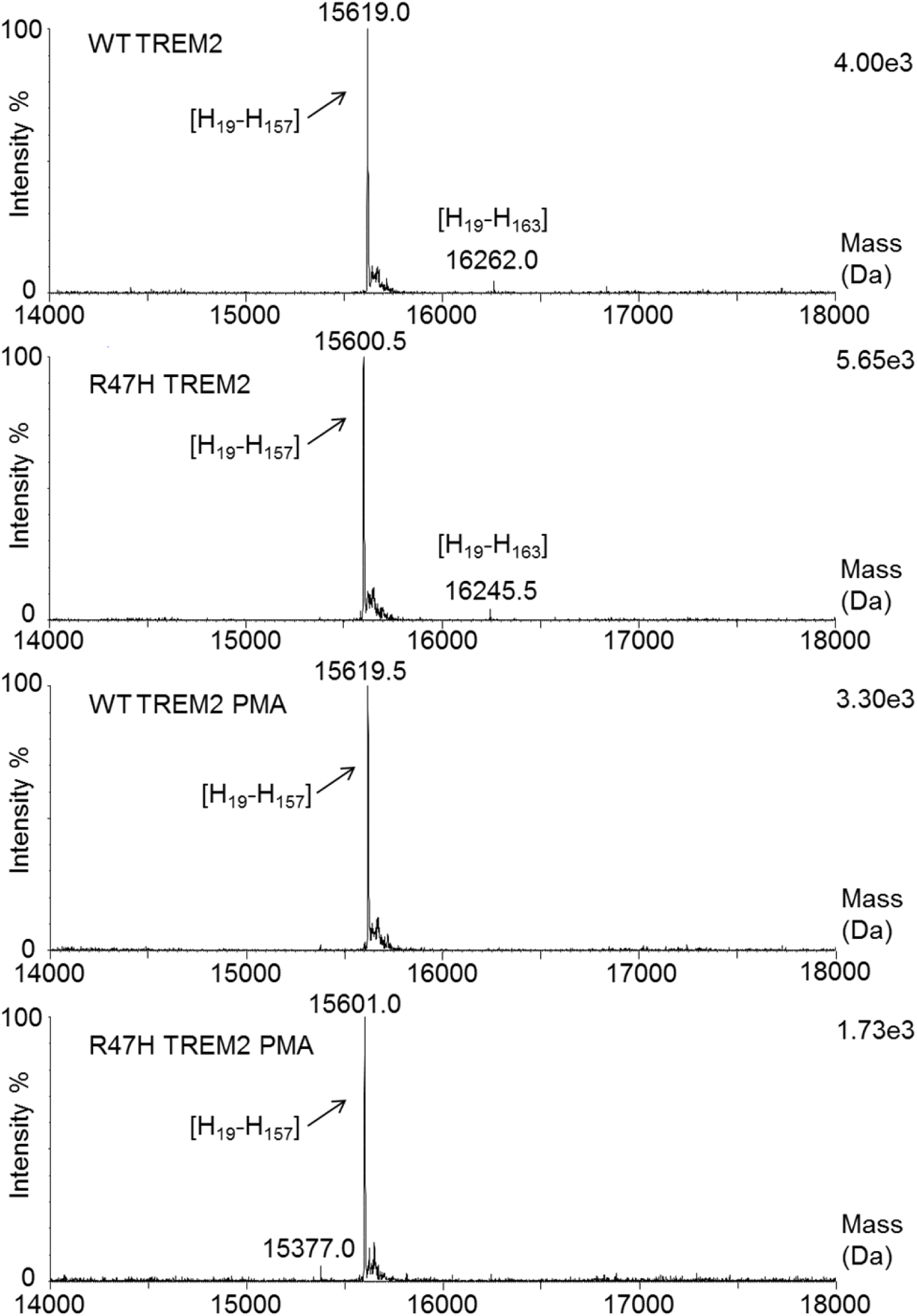
Identification of shed TREM2 ectodomain from HEK-FT cells transiently transfected with WT or mutant R47H hTREM2 and hDAP12. Deconvoluted mass spectra of 4 mass spectra: the shed hTREM2 [19-157]is clearly present in all four cell supernatant extracts. The cell supernatants had been treated with PNGase-F and Sialidase A after affinity purification, but not reduced.

### TREM2 is not O-glycosylated at positions close to the cleavage site

As post-translational modifications can effect sheddase activity (26,27), we sought to identify how glycosylation might effect TREM2 shedding. Two putative O-glycosylation sites exist close to the identified H157 cleavage site of the TREM2 ectodomain: S160 and S168. Using C-terminally his-tagged hTREM2 and mapping glycosylation sites by mass spectrometry, we could show that hTREM2 displays O-glycosylation at T171 and/or S172 (figure 7), however we could not detect any O-glycosylation at S160 or S168.

**FIGURE 7.**
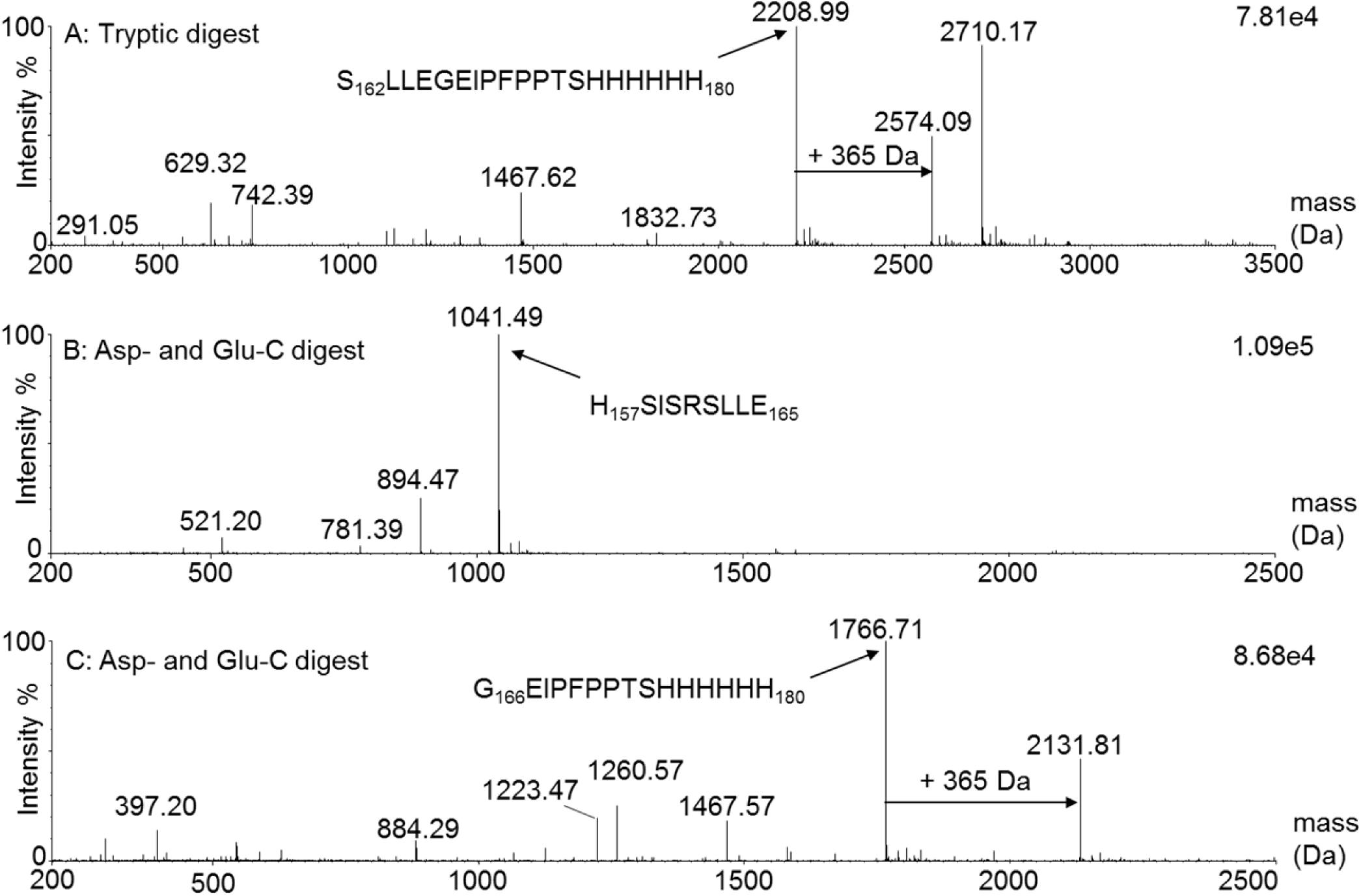
Determination of O-glycosylation site(s) within TREM2 stalk region. TREM 2-His was first treated with Sialidase A, then reduced, alkylated, subsequently treated by PNGase-F. The resulting sample was then either digested by trypsin or by Asp-and Glu-C enzyme. The digests were analyzed by LC-MS^E^. Panel A: Deconvoluted mass spectrum, combined scans: 1926:2097. Panel B: Deconvoluted mass spectrum, combined scans: 1566:1584). Panel C: Deconvoluted mass spectrum, combined scans: 1373:1477).

## Discussion

This study first investigated the contribution of ADAM10 and ADAM17 to shedding of TREM2. The pharmacological and genetic approaches used show unambiguously that ADAM17 is the pivotal protease contributing to this process. Enzyme selectivity of the ADAM inhibitors applied (GI254023 (22) and DPC333(21)) was determined in an *in vitro* cleavage assay and both inhibitors were used over a broad concentration range to investigate effects on TREM2 cell surface expression in CHO-hDAP12-hTREM2 cells and human M2A macrophages. To rule out that the pharmacological approach was not confounded by unspecific inhibition of other enzymes, these findings were corroborated by CRISPR/CAS9 knockout of ADAM10 or ADAM17 in the human monocytic THP-1 cell line. The knockout experiments confirmed that ADAM17 deficiency increased surface TREM2 and reduced sTREM2. The data presented here suggest that under steady state conditions TREM2 is cleaved mainly by ADAM17, but after activation of ADAMs by PMA, additional shedding mechanisms might come into play. Recent literature data indicated that under steady state conditions ADAM10 mediates generation of sTREM2 (14). The main difference between that study and the results presented here is the use of HEK-Flp-In cells which lack co-expression of the signaling adaptor protein DAP12 together with TREM2. It might be that TREM2 after expression in a recombinant system in the absence of DAP12 has increased susceptibility to ADAM10 cleavage. Interestingly DAP12 has a 14 AA long extracellular domain (See (28) for review). The close proximity of the extracellular domain of DAP12 to the TREM2 cleavage site could suggest an interplay between the extracellular portions TREM2 and DAP12 that could regulate activation and shedding.

In addition to the investigation of constitutive shedding, we used PMA to enhance shedding of TREM2 from the cell surface. Shedding observed under these conditions in the ADAM17 deficient THP1 cells could be attributed to ADAM10 activity, but might also involve other mechanisms. E.g. TREM2 ectodomain might be cleaved intracellularly during its transport from the ER to the plasma membrane, allowing for TREM2 secretion into the medium.

Recent studies have elucidated how PMA treatment and changes in sheddase activity are connected: PMA-triggered signaling cascades activate scramblases that enhance translocation of phosphatidylserine (PS) to the outer leaflet of the plasma membrane (29). Here PS binds to a cationic motif in the membrane proximal domain of ADAM17 enabling the protease to execute its sheddase function (23). These experiments focused on ADAM17 and further studies are required to elucidate whether any of these findings extend to ADAM10, the closest homolog to ADAM17 within the ADAM family.

It is of note that PS has also been described as TREM2 ligand (15,30-32). Phosphatidylserine belongs to a range of membrane phospholipids which are exposed by damaged neurons and glial cells or released by damaged myelin. Further, negatively charged phospholipids like PS have been shown to associate with Aβ in lipid membranes (33,34). In all these processes, PS acts as ligand for TREM2, and could at the same time facilitate its shedding.

Our mutational analysis identified two AA stretches within the stalk region of TREM2 which are important for TREM2 shedding. The membrane distal area comprises the AA of the cleavage site. Interestingly, exchanging 5 AA close to the plasma membrane (mutant T2-WFRW) also stabilized TREM2 at the cell surface. Although this has not been investigated for TREM2, co-clustering of ADAM17 with its substrate L-selectin (35) has been described and this part of the stalk region might contribute to this process enabling binding of ADAM17 to its substrate before cleavage is initiated upon activation of the proteolytic activity of ADAM17 (e.g. by PS).

Additional experiments identified the precise cleavage site for the TREM2 ectodomain. Three complementary approaches were applied, and all show the same cleavage site. First, peptides from the stalk region were subjected to *in vitro* cleavage with recombinantly expressed ADAM10 and ADAM17. Second, we determined the C-terminus from tryptic peptides of the TREM2 ectodomain purified from supernatant of HEK-FT cells recombinantly expressing hTREM2 and hDAP12. Third, we verified the size of full length TREM2 ectodomain purified from cellular supernatant. While the distance of the cleavage site from the plasma membrane (17 AA) is at the upper end when compared to most known ADAM17 substrates (12-16 AA, (36,37)), the sequence (P2:V, P1:H, P1’:S, P2’:I) is quite unique compared to known ADAM17 or ADAM10 substrates (38-41). However, if cleavage sites for both ADAM17 and ADAM10 are compiled, (40) arginine at P1 is quite common. Histidine at this position in TREM2 also carries a basic side chain with a positive charge. It is noteworthy that there was no shift of cleavage to another side within the peptide. This supports the observation that shedding seems to be confined to a region close to the plasma membrane and no shift of cleavage to a secondary site occurs. This is line with earlier findings suggesting that the position of the site relative to the transmembrane region and the first globular part of the protein is as important as the AA sequence of the cleavage site (36,42,43).

The R47H variant confers a significantly increased risk to develop LOAD (11,12). In contrast to some of the polymorphisms that confer increased risk for FTD like T66M (11,14,44,45) and Y38C (11), which reduce cell surface expression of TREM2, the R47H mutation does not affect expression levels but rather functionally impairs TREM2 (8,9,14-16). Data presented here indicate that R47H mutation does not affect cleavage site, i.e. the size of soluble, shed TREM2 ectodomain. However, these experiments do not allow to conclude whether extent of shedding is changed, i.e. the amount of sTREM2 is altered.

Ectodomain shedding has been described to be influenced by O-glycosylation at serine or threonine residues which are within ± 4 residues of the processing site (26,27). We investigated the presence of O-glycosylation close to the cleavage site. Our results indicate that within the stalk region TREM2 is only O-glycosylated at T171 and/or S172, but not at S168 or S160. Most likely this O-glycosylation site is too remote from the processing site to influence cleavage. These results are limited by the fact that the hTREM2 protein used for these studies was not shed from the cell surface but secreted through the secretory pathway in the expression system used for the production of the recombinant protein.

In the recent years a wealth of TREM2 variants have been identified that influence susceptibility for certain neurodegenerative diseases (See (46) for a recent summary). Most interestingly, one of these mutations H157Y (rs2234255, (32,47-49)) is located precisely at the identified TREM2 cleavage site, and increases the propensity to develop AD. *In vitro* data show that ADAM10/17 has an increased preference for tyrosine in the P1’ position (40). One could therefore speculate that this mutation may lead to enhanced constitutive shedding of TREM2 from the cell surface by ADAM10/17, and provide a mechanistic link between TREM2 shedding and development of AD, but exactly how this mutation affects shedding of TREM2 is currently unknown.

A fascinating question that arises from these results is whether sTREM2 has a physiological role. In initial phases of host defense or sterile inflammation, robust inflammation is advantageous for pathogen neutralization or removal of damaged tissue followed by the subsequent resolution response (50). Upon resolution of inflammation, ADAM17 activity would diminish (51,52) and the resulting increase in TREM2 would promote both resolution and phagocytosis. At the same time production of other pro-inflammatory cytokines like TNFα will become reduced.

## Experimental procedures

### Compounds

GI254023 ((2R,3S)-3-(formyl-hydroxyamino)-2-(3-phenyl-1-propyl) butanoic acid [(1S)-2,2-dimethyl-1-methylcarbamoyl-1-propyl] amide) was synthesized as described in (22)) (Blood. 2003;102:1186-1195). DPC333 ((2R)-2-((3R)-3-amino-3{4-[2-methyl-4-quinolinyl) methoxy] phenyl}-2-oxopyrrolidinyl)-N-hydroxy-4-methylpentanamide)) was synthesized as described in (21).

### Cell Culture

THP1 cells stably coexpressing Cas9 and a blasticidin resistance gene delivered by lentivirus were cultured in RPMI medium containing 10% FBS, 1% L-glutamine, 1% pen/strep, and 10?g/ml of blasticidin (Thermo Fisher Scientific). The cells were cultured at 37°C in 5% CO2 atmosphere.

### Generation of ADAM17 and ADAM10 Knockout lines

THP1-Cas9 cells were infected with lentiviruses expressing the puromycin resistance gene and sgRNAs (for vector design, see (53)) targeting either ADAM10 (GTAATGTGAGAGACTTTGGG) or ADAM17 (CCGAAGCCCGGGTCATCCGG). Lentiviral packaging was carried out in HEK293T cells as described in. 30 μL of lentiviral sgRNA supernatant was added to 1x10^6^ THP1-Cas9 cells in 2ml medium containing 5 μg/ml polybrene (Sigma) and spun at 300g for 90 min. in a 6-well plate. After 24 hours the cells were spun down and resuspended in fresh culture medium containing 1.5 μg/mL puromycin. After 4 weeks of weekly media changes, genomic DNA was isolated using a Quick-gDNA miniprep kit (Zymo Research) to assess the insertions and deletions (indels) present in the pool by next-generation sequencing (NGS). To isolate clones containing only frameshift indels, cells were plated at limiting dilution into 96-well plates. Upon expansion of the clones, they were assayed by NGS and clones containing only frameshift alleles were selected for downstream assays.

### NGS indel analysis

To prepare engineered cells for NGS, each target was amplified using locus-specific primers. Two rounds of PCR were performed. The first round utilized locus-specific primers to amplify the edited region. The primers for ADAM10 were ATTAGACAATACTTACTGGGGATCC and GGAAGCTCTGGAGGAATATGTG, and the primers for ADAM17 were CCCCCAAACACCTGATAGAC and CCAGAGAGGTGGAGTCGGTA. The product formed during the first round was then used as a template for a second round of PCR to add dual indices compatible with the Illumina system. The Illumina Nextera Adapter sequences for both target regions were TCGTCGGCAGCGTCAGATGTGTATAAGAGACAG and GTCTCGTGGGCTCGGAGATGTGTATAAGAG ACAG. Libraries were quantitated by qRT-PCR and subsequently sequenced on the Illumina MiSeq system. For sequence analysis, raw reads were aligned to a reference sequence, then tallied based on genotype. Finally, tallied genotypes were binned into one of three categories: wild-type, in-frame, and frameshift.

### Cell surface expression of mutated TREM2 in HEK293 cells

Human TREM2 mutants were generated using the QuikChange Site-directed Mutagenesis Kit (Stratagene) and confirmed by sequence analysis. Transfections of complementary DNA (cDNA) constructs were carried out in a 1:1 ratio of hDAP12 and TREM2 in HEK-FT cells using Lipofectamine LTX reagent (Thermofischer) according to the manufacturer’s recommendations. 48 h after transfection cells were treated for 30 min with 50 ng/ml PMA or 0.05% DMSO and detached with a cell detachment solution Accutase (Sigma) and stained with goat anti human TREM2 antibody AF1828 (R&D Systems) or isotype control followed by incubation with the secondary Alexa Fluor 488 conjugated antibody (Molecular Probes). Acquisition was performed using a BD FACSCanto II (BD biosciences).

### TREM2 expression in human macrophages and CHO cells

– CHO cells were transfected to co-express hDAP12 and hTREM2 using Lipofectamine LTX reagent (Thermofischer) according to the manufacturer’s recommendations. One positive clone was selected and designated CHO-hDAP12-hTREM2. Human M2A macrophages were obtained from buffy coat using a negative isolation kit for monocytes (Stem cell technologies) and differentiated for 5 days in RPMI1640 medium with Glutamax (Gibco) supplemented with 10% FBS (Gibco), PenStrep (Gibco), 1 % Sodium Pyruvate (Gibco), 0.025 M HEPES Buffer (Gibco), 0.05 mM β-mercaptoethanol (Gibco), M-CSF (40 ng/ml) and IL-4 (50 ng/ml). Cell surface TREM2 was detected by FACS as described above.

### Life cell imaging

Human M2A macrophages or CHO-hDAP12-hTREM2 were seeded on 384 well plates (Greiner) and treated with ADAM inhibitors DPC333 or GI254023 at concentrations indicated in the figures. 16 h later cells were treated for 30 minutes with PMA (50 ng/ml) or 0.1 % DMSO for 30 minutes. Plates were put on ice and stained with goat anti human TREM2 antibody AF1828 (R&D Systems) or isotype control and Hoechst stain followed by incubation with the secondary Alexa Fluor 488 conjugated anti goat antibody (Molecular Probes). Images were acquired using an InCell2000 analyzer (GE Healthcare). For image quantification the free open source software CellProfiler was applied. Reporter gene assay in BWZ cells – BWZ thymoma reporter cells, which express lacZ under control of the promoter for nuclear factor of activated T cells (NFAT, Hsieh 2009) were transfected to co-express mDAP12 and WT hTREM2 or T2double TREM2. Cells were seeded in RPMI without phenol red supplemented with 2% FBS and 1% non-essential amino acids on high binding microtiter plates (Greiner) pre-coated with rat anti-mouse/human TREM2 mAb (R&D, MAB17291) or isotype control. Cell culture was continued for 16 h. Reporter gene activity was assessed with the Beta-glo assay system (Promega) according to the manufacturer’s recommendations using an Envision 2104 multilabel reader (Perkin Elmer).

### Immuno-purification

Shed TREM2 was purified from cell supernatant through microscale immuno-purification. This was performed on the MEA platform (PhyNexus) and Streptavidin coated tips (PTR 92-05-05, Phynexus) have been used. First the tips are equilibrated with PBS, then 200 μL of biotinylated anti-TREM2 antibody (0.55 pmol/μL, BAF1828 from R&D System), is loaded onto the streptavidin μcolumn (5 μL bed volume) at a speed of.25 mL/min and 8 passages. After a wash with PBS, the shed TREM2 is captured from the cell supernatant (200 μL) at a speed of 25 mL/min and 12 passages. It is followed by PBS wash and elution by 0.1M Glycine pH 2.5 (2 x 4 passages) for a final volume of 2x60 μL. The latter solution is neutralized with the addition of 1 M Tris-HCl pH 10 (5 μl), then it was dried (Speedvac) and rehydrated with 8M urea (5 μL, Fluka) and 0.4M NH4CO3 (30 μL, Fluka). The sample is then reduced (2 μL of 1M DTT, 30 min at 50°C), alkylated (6 μl of 1M IAA (Sigma), 30min at RT in dark) and the reaction was terminated with the addition of 1M DTT (2 μL) and 0.4M NH4CO3 (30 μl). The resulting sample is either digested by Trypsin or Asp-/Glu-C enzyme (+1 μl of Trypsin (Promega) or Asp-/Glu C (Roche), 1μg/μl, pH 8, and overnight incubation at 37°C). The digested sample is finally acidified with HCOOH (1 μL, Fluka) and 25 μL of the resulting digest were injected onto the LC-MS platform.

### Mass spectrometry

Identification of the cleavage site by peptide mapping: the LC-MS^E^ analyses were performed using a SYNAPT G2S QTOF mass spectrometer (WATERS) coupled with an UPLC (ACQUITY I class, WATERS). A BEH C18 UPLC column (1.7 μm, 1×100 mm, WATERS) was used for peptide separation. An elution gradient with mobile phase A (0.1% HCOOH in water) and mobile phase B (0.1% HCOOH in acetonitrile) was generated using the following program: 1) isocratic at 2% B for 3 min; 2) linear gradient from 2 to 30% B from 3 to 90 min; 3) linear gradient from 30 to 100% B from 90 to 95 min; 4) isocratic at 100% B from 95 to 105 min; 5) linear gradient from 100 to 2% B from 105 to 105.5 min; and finally 6) isocratic at 2% B from 105.5 to 120 min. The mass spectrometer was working in positive resolution mode with automatic mass correction through a lockspray system (P_14_R peptide, m/z 767.433, infused at 250 fmol at 5 μL/min, switch frequency was every 20 sec for 0.5 sec per scan, 3 scans averaged). 2 MS traces were acquired, one MS and one for MS^E^. Both were acquired in mass range m/z 50 – 2000, scan time 0.5 sec, 3 kV capillary voltage, 40 V cone voltage. In MS^E^ mode the trap voltage was ramped in each scan from 20 to 40 V. In addition, a UV trace was acquired at a wavelength of 214 nm.

### Identification of the cleavage site by intact mass measurement

The LC-MS analyses were performed using a SYNAPT G1 QTOF mass spectrometer (WATERS) coupled with an UPLC (ACQUITY I class, WATERS). A BEH C4 UPLC column (1.7 μm, 1×100 mm, WATERS) was used for protein separation. An elution gradient with mobile phase A (0.1% HCOOH in water) and mobile phase B (0.1% HCOOH in acetonitrile) was generated using the following program: 1) isocratic at 5% B for 1.5 min; 2) linear gradient from 5 to 25% B from 1.5 to 2 min; 3) linear gradient from 25 to 35% B from 2 to 12 min; 4) linear gradient from 35 to 95% B from 12 to 13 min; 5) isocratic at 95% B from 13 to 15 min; 5) linear gradient from 95 to 5% B from 15 to 15.5 min; and finally 6) isocratic at 5% B from 15.5 to 20 min. The mass spectrometer was working in positive resolution mode and calibrated with NaI 2mg/mL. MS trace was acquired in mass range m/z 600 – 4500, scan time 0.5 sec, 3 kV capillary voltage, 40 V cone voltage, desolvation temperature 200 °C, cone gas flow 50 L/h. In addition, a UV trace was acquired at a wavelength of 214 nm.

### Identification of O-linked glycosylation

The LC-MS^E^ analyses were performed using a QTOF Premier mass spectrometer (WATERS) coupled with an UPLC (ACQUITY H class, WATERS). A BEH C18 UPLC column (1.7 μm, 1×100 mm, WATERS) was used for peptide separation. An elution gradient with mobile phase A (0.1% HCOOH in 98% water and 2 % acetonitrile) and mobile phase B (0.1% HCOOH in acetonitrile) was generated using the following program: 1) isocratic at 2% B for 3 min; 2) linear gradient from 2 to 30% B from 3 to 90 min; 3) linear gradient from 30 to 100% B from 90 to 95 min; 4) isocratic at 100% B from 95 to 105 min; 5) linear gradient from 100 to 2% B from 105 to 105.5 min; and finally 6) isocratic at 2% B from 105.5 to 120 min. The mass spectrometer was working in positive normal mode with a lockspray system (P_14_R peptide, infused at 1 pmol at 10 μL/min, switch frequency was every 20 sec for 0.5 sec per scan, 3 scans averaged). Mass correction was performed by application of a lockmass (2+, 767.433 Da) during data processing with PLGS (WATERS). 2 MS traces were acquired, one MS and one for MS^E^. Both were acquired in mass range m/z 50 – 2000, scan time 0.5 sec, 3 kV capillary voltage, 40 V cone voltage. In MS^E^ mode the trap voltage was ramped in each scan from 20 to 40 V. In addition, a UV trace was acquired at a wavelength of 214 nm.

### Statistical analysis

Statistical analysis was performed using Prism software (GraphPad, San Diego, CA) using ANOVA and student’s t test where appropriate. A p value of < 0.05 was considered significant.

## Author Contributions

FD, NU and SP conceived and coordinated the study and wrote the paper. FD, BC, EG, KL, BI, B-LE, WKA, HDJ, YC and JS performed experiments, analyzed and interpreted data. WK, MA, FC, NMC, HCL, KA, PS, GN and AR contributed to the conception of the study and the interpretation of data. All authors examined the results and approved the final version of the manuscript.

## Background

TREM2 is associated with neurodegenerative diseases and its ectodomain is shed during inflammatory conditions.

## Results

ADAM17 is identified as main shedding enzyme for TREM2 ectodomain and cleavage occurs at H-S (AA 157-158) within the stalk region. R47H mutation does not affect shedding.

## Conclusion

Activity of ADAM17 during disease regulates sTREM2 production and determines TREM2 cell surface expression.

## Significance

Sheddases influence activity of microglia and macrophages during inflammation.

## The abbreviations used are

TREM2: Triggering receptor expressed on myeloid cells
AA: amino acids
DAP12: DNAX-activating protein 12
ADAM1: a disintegrin and metalloproteinase domain containing protein
TACE: TNFα converting enzyme

## Supplementary figures

**S-FIGURE 1:**
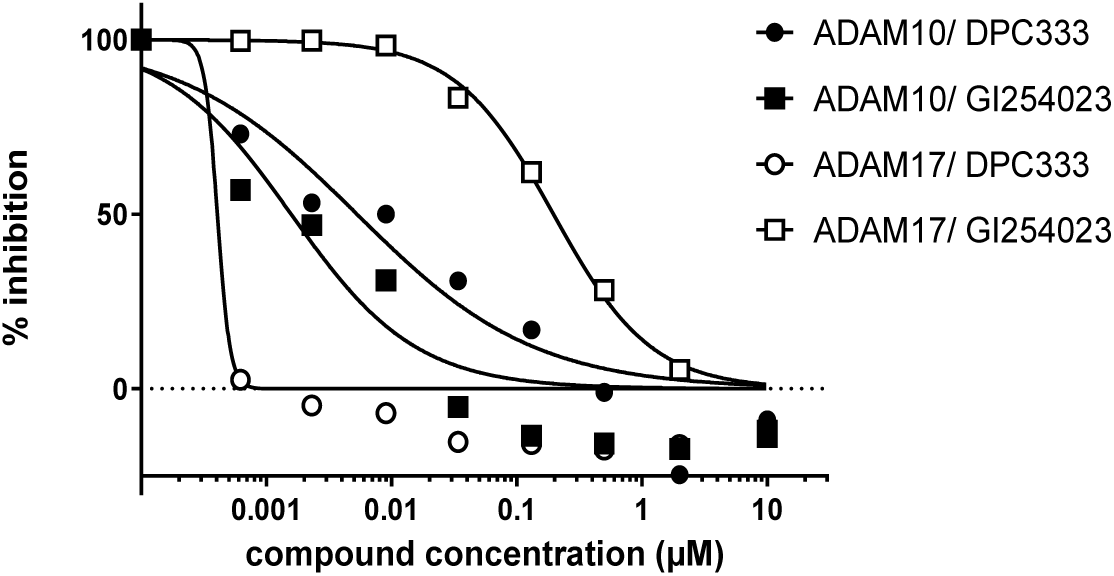
*In vitro* selectivity of DPC333 and GI254023 at human ADAM10 and ADAM17. Inhibition of recombinant ADAM10 and ADAM17 by DPC333 and GI254023 *in vitro* In Hepes buffer pH 7.5.

**S-FIGURE 2:**
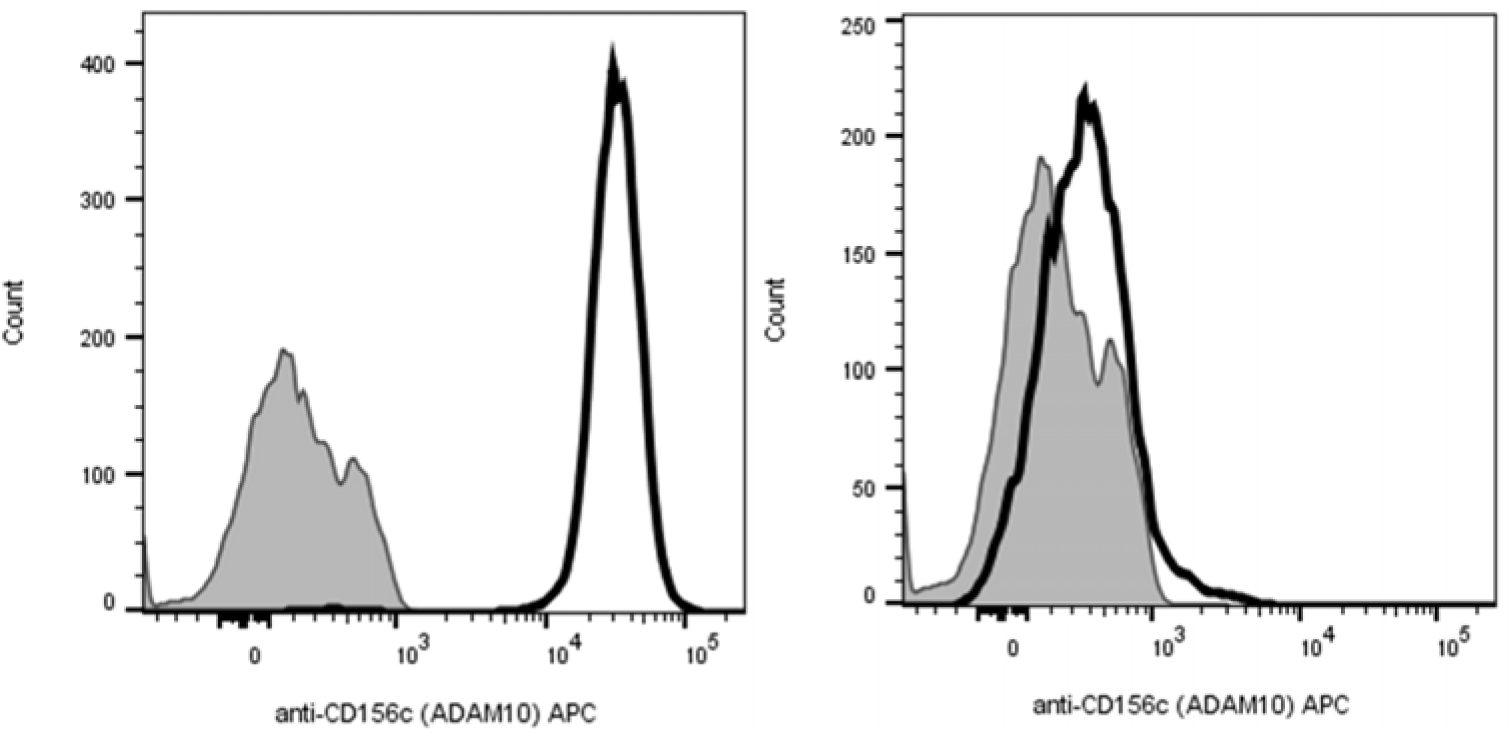
Lack of ADAM10 expression in THP1 ADAM10 H4 CRISPR clone. FACS analysis of ADAM10 cell surface expression in THP1 CRISPR cell clones. Left panel control clone CtrlgRNA, right panel AD10 H4 clone.

**S-FIGURE 3:**
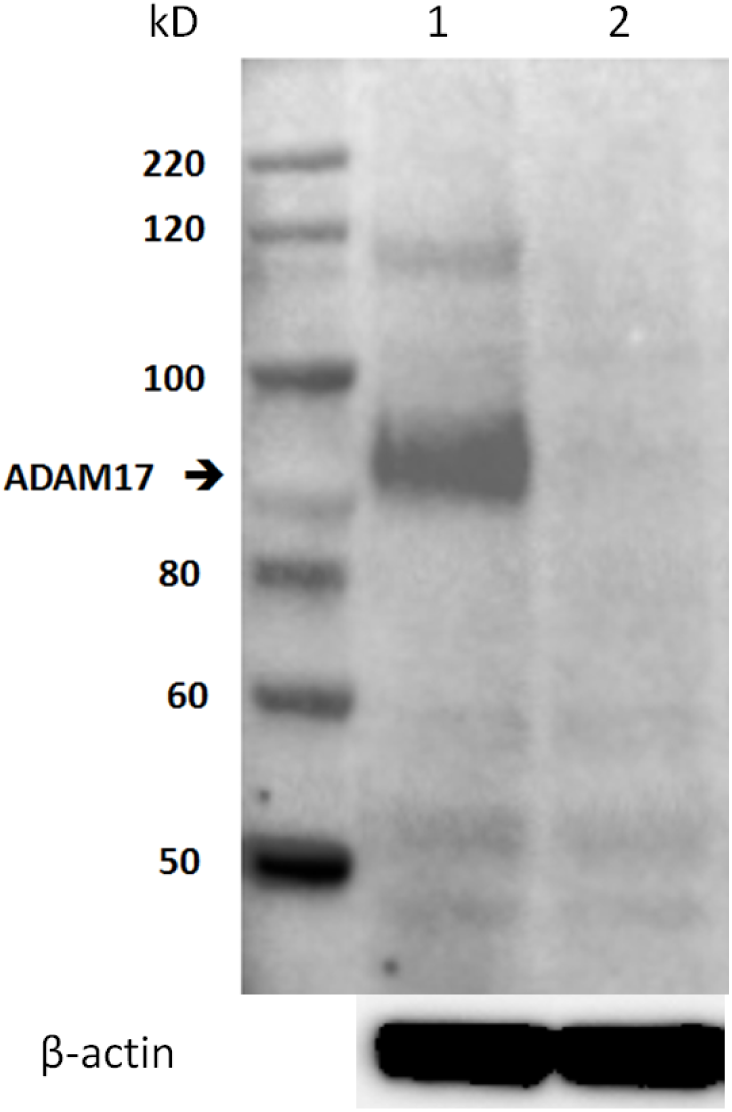
Lack of ADAM17 expression in THP1 ADAM17 G12 CISPR clone. Western analysis of ADAM17 expression in THP1 control clone CtrlgRNA (lane 1) and ADAM17 AD17 G12 CRISPR cells (lane 2).

**S-FIGURE 4:**
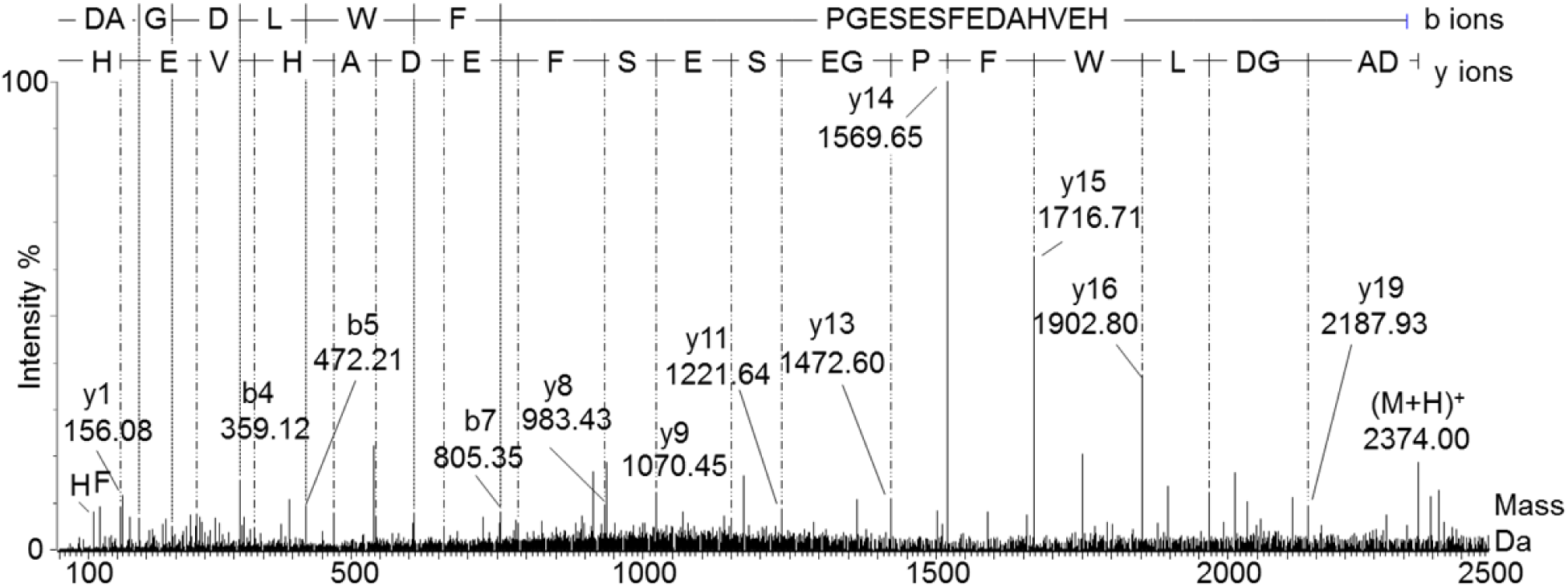
Example of MS^E^ spectrum of peptide D_137_-H_157_. Deconvoluted MS^E^ spectrum of peptide D_137_-H_157_. Top panel indicates which b and y fragment ions are identified. This mass spectrum is from the tryptic digest of TREM2 R47H PMA.

